# The unjamming transition is distinct from the epithelial-to-mesenchymal transition

**DOI:** 10.1101/665018

**Authors:** Jennifer A. Mitchel, Amit Das, Michael J. O’Sullivan, Ian T. Stancil, Stephen J. DeCamp, Stephan Koehler, James P. Butler, Jeffrey J. Fredberg, M. Angela Nieto, Dapeng Bi, Jin-Ah Park

## Abstract

Every organ surface and body cavity is lined by a confluent collective of epithelial cells. In homeostatic circumstances the epithelial collective remains effectively solid-like and sedentary. But during morphogenesis, remodeling or repair, as well as during malignant invasion or metastasis, the epithelial collective becomes fluid-like and migratory^1–4^. This conversion from sedentary to migratory behavior has traditionally been understood as a manifestation of the epithelial-to-mesenchymal transition (EMT) or the partial EMT (pEMT)^5–8^. However, in certain contexts this conversion has been attributed to the recently discovered unjamming transition (UJT), in which epithelial cells move collectively and cooperatively^9–11^. UJT and pEMT share certain aspects of collective cellular migration, but the extent to which these processes are distinct, overlapping or perhaps even identical has remained undefined. Using the confluent layer of well-differentiated primary human bronchial epithelial (HBE) cells, here we triggered UJT by exposing the sedentary layer to mechanical compression^9–12^. Cells thereafter migrated cooperatively, aligned into packs locally, and elongated systematically. Nevertheless, cell-cell junctions, apico-basal polarity, and barrier function remained intact in response, and mesenchymal markers remained unapparent. As such, pEMT was not evident. When we triggered pEMT and associated cellular migration by exposing the sedentary layer to TGF-β1, metrics of UJT versus pEMT diverged. To account for these striking physical observations a new mathematical model attributes the effects of pEMT mainly to diminished junctional tension but attributes those of UJT mainly to augmented cellular propulsion. Together, these findings establish that UJT is sufficient to account for vigorous epithelial layer migration even in the absence of pEMT. Distinct gateways to cellular migration therefore become apparent – UJT as it might apply to migration of epithelial sheets, and EMT/pEMT as it might apply to migration of mesenchymal cells on a solitary or collective basis, activated during development, remodeling, repair or tumor invasion. Through the actions of UJT and pEMT working independently, sequentially, or interactively, living tissue is therefore seen to comprise an active engineering material whose modules for plasticity, self-repair and regeneration, are far richer than had been previously appreciated.

---

Since its discovery in 1982, the epithelial-to-mesenchymal transition (EMT) has been intensively studied and well-characterized^6, 13, 14^. EMT is marked by progressive loss of epithelial character, including disrupted apico-basal polarity, disassembled cell-cell junctions, and impaired epithelial layer integrity and barrier function. This loss of epithelial character is accompanied by progressive gain of mesenchymal character, including gain of front-back polarity, activation of EMT-inducing transcription factors, and expression of mesenchymal markers^15^. In this process each epithelial cell tends to free itself from adhesions to immediate neighbors, and thereby can acquire migratory capacity and invasiveness. It has been suggested that the epithelial-mesenchymal axis is flanked at its extremes by unequivocal epithelial versus mesenchymal phenotypes separated by a continuous spectrum of hybrid epithelial/mesenchymal (E/M) or partial EMT (pEMT) phenotypes^5, 8^. Although such a one-dimensional spectrum of states has been regarded by some as being overly simplistic^5, 16^, it is widely agreed that pEMT allows cell migration without full mesenchymal individualization^17–19^. During pEMT, cells coordinate with their neighbors through intermediate degrees of junctional integrity coupled with partial loss of apical-basal polarity and acquisition of graded degrees of front-back polarity and migratory capacity^5, 8^. Moreover, EMT/pEMT is associated with the cells of highly aggressive tumors, endows cancer cells with stemness and resistance to cytotoxic anticancer drugs, and may be required in the fibrotic response^20, 21^. In development^22–25^, wound healing^26, 27^, fibrosis^28^ and cancer^29–33^, EMT/pEMT has provided a well-accepted framework for understanding collective migration, and in many contexts has been argued to be necessary^2, 17, 21, 27, 34, 35^.

By contrast with EMT, the unjamming transition (UJT) in epithelial tissues is newly discovered and remains poorly understood^9–11, 36–44^. During UJT the epithelial collective transitions from a jammed phase wherein cells remain virtually locked in place, as if the cellular collective were frozen and solid-like, toward an unjammed phase wherein cells often migrate in cooperative multicellular packs and swirls reminiscent of fluid flow. In both the solid-like jammed phase and the fluid-like unjammed phase, the epithelial collective retains an amorphous disordered structure. In the jammed phase, the motion of each individual cell tends to be caged by its nearest neighbors, but as the system progressively unjams and transitions to a fluid-like phase, local rearrangements amongst neighboring cells become increasingly possible, and tend to be cooperative, intermittent and heterogeneous^41, 45–47^. While poorly understood, cellular jamming and unjamming have been identified in epithelial systems *in vitro*^9, 11, 38, 41, 43, 44^, in developmental systems in *vivo*^11, 37^, and have been linked to the pathobiology of asthma^9–11^ and cancer^40, 42^.

Despite strong evidence implicating both pEMT and UJT in the solid-fluid transition of a cellular collective and the resulting collective migration of cells of epithelial origin^9–11, 34^, the relationship between these transitions remains undefined. For example, it is unclear if UJT entails elements of the pEMT program. The converse is also in question. As such, we do not yet know if the structural, dynamical, and molecular features of these solid-fluid transitions might be identical, overlapping, or entirely distinct. To discriminate among these possibilities, here we examine mature, well-differentiated primary human bronchial epithelial (HBE) cells grown in air-liquid interface (ALI) culture; this model system is known to recapitulate the cellular constituency and architecture of intact human airway epithelia^48–50^. To induce UJT we exposed the cell layer to mechanical compression (30 cm H_2_O) mimicking the physical forces experienced by the epithelial layer during asthmatic bronchoconstriction^12, 51–58^. To induce pEMT we exposed the cell layer to TGF-β1 (10 ng/ml), a well-known EMT-inducing agent^21, 59^.

## Cellular dynamics and structure: UJT versus pEMT diverge

In a sedentary confluent epithelial layer, initiation of either UJT or pEMT results in collective migration^8–11, 18, 34^. While the precise dynamic and structural characteristics of the HBE layer undergoing pEMT have not been previously explored, UJT is known to be marked by the onset of stochastic but cooperative migratory dynamics together with systematic elongation of cell shapes^9–11, 44, 60–62^.

### Dynamics

We quantified migratory dynamics using average cell speed and effective diffusivity (D_eff_)^9,62^. Control HBE cells were essentially stationary, as if frozen in place, exhibiting only occasional small local motions which were insufficient for cells to uncage or perform local rearrangements with their immediate neighbors (Fig. 1a-c). We refer to this as kinetic arrest or, equivalently, the jammed phase. Following exposure to mechanical compression, however, these cells underwent UJT and became migratory^9–11^, with both average speed and effective diffusivity increasing substantially over time and maintained to at least 72 hrs following compression (Fig. 1a-c, Extended Data Table 1). Following exposure to TGF-β1, jammed cells underwent pEMT^21, 59^, as documented in greater detail below (Fig. 2, Extended Data Figs. 1 and 2). Up to 24 hrs later these cells migrated with average speeds comparable to cells following compression (Fig. 1a-c). However, as pEMT progressed beyond 24 hrs cellular motions slowed to the baseline levels, indicating return to kinetic arrest and a jammed phase (Fig. 1a-c, Extended Data Table 1).

**Figure 1.**
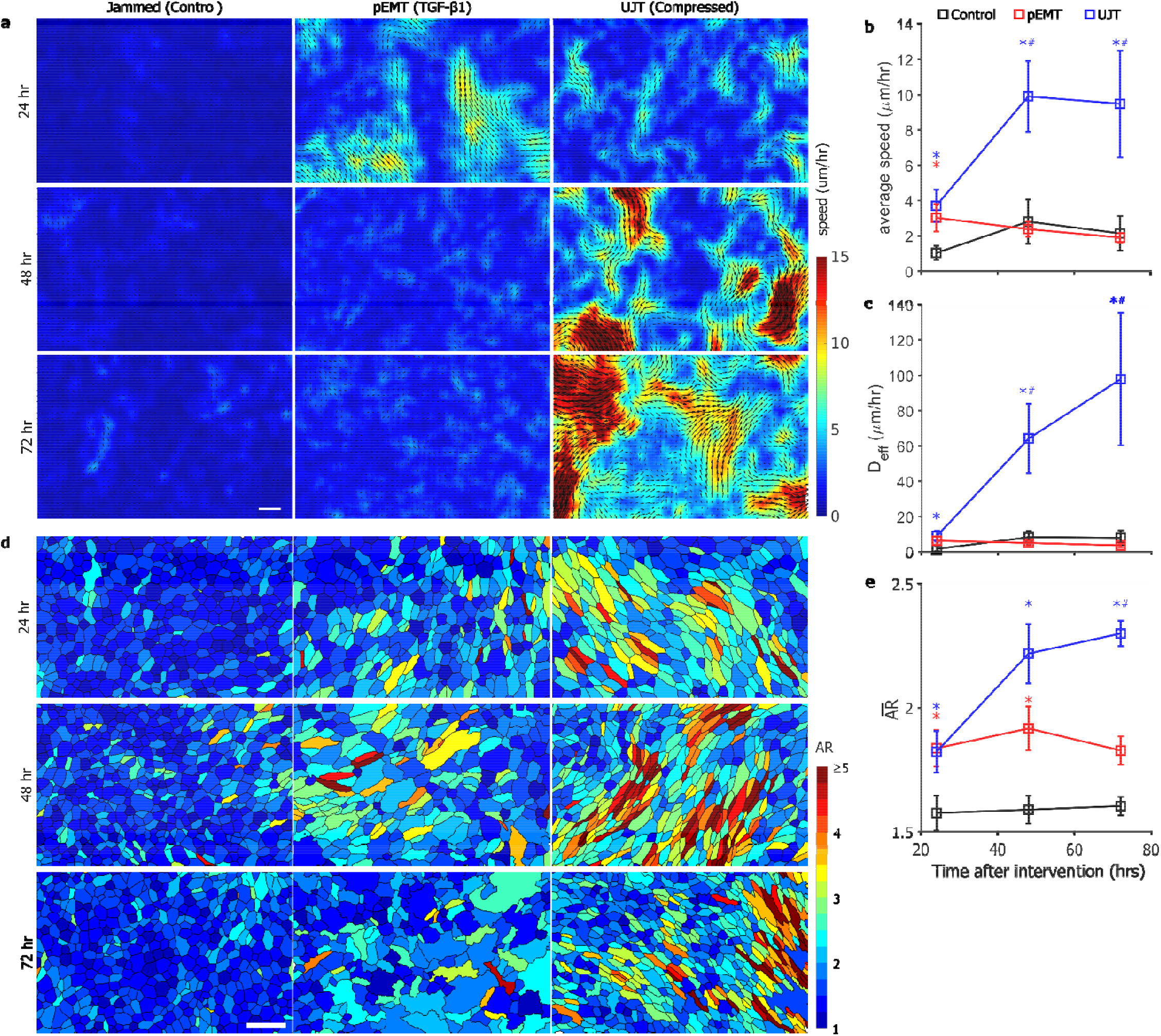
In both dynamics and structure, pEMT and UJT diverge over time. In well-differentiated HBE cells, pEMT was induced by TGF-β1 (10 ng/ml) and UJT was induced by compression^9, 12, 56, 97^. <u>Dynamics:</u> Representative speed maps (obtained using optical flow, see methods) (**a**), average speed (**b**), and effective diffusivity (**c**) for jammed, pEMT and UJT at 24, 48, and 72 hrs (6-12 fields of view per condition and timepoint, n=4 donors). Jammed cells remain essentially stationary over all times. Cells undergoing pEMT migrate moderately at 24 hrs but return to baseline by 48 hrs. Cells undergoing UJT display progressively increasing migratory speed and diffusivity over time. Scale bar in **a** is 100 μm. <u>Structure:</u> Individual cells color-coded by aspect ratio (**d**) and quantified by mean AR (**e**) for jammed, pEMT, and UJT at 24, 48 and 72 hrs from at least two fields of view per condition and timepoint for each of n=3 donors. Elevated cellular AR represents a structural signature of the unjamming transition^9, 11^. Scale bar in **d** is 50 μm. See also Extended Data Table 1. *p<0.05, vs. control; ^#^p<0.05, UJT vs. pEMT, color coded according to which sample is referenced.

**Figure 2.**
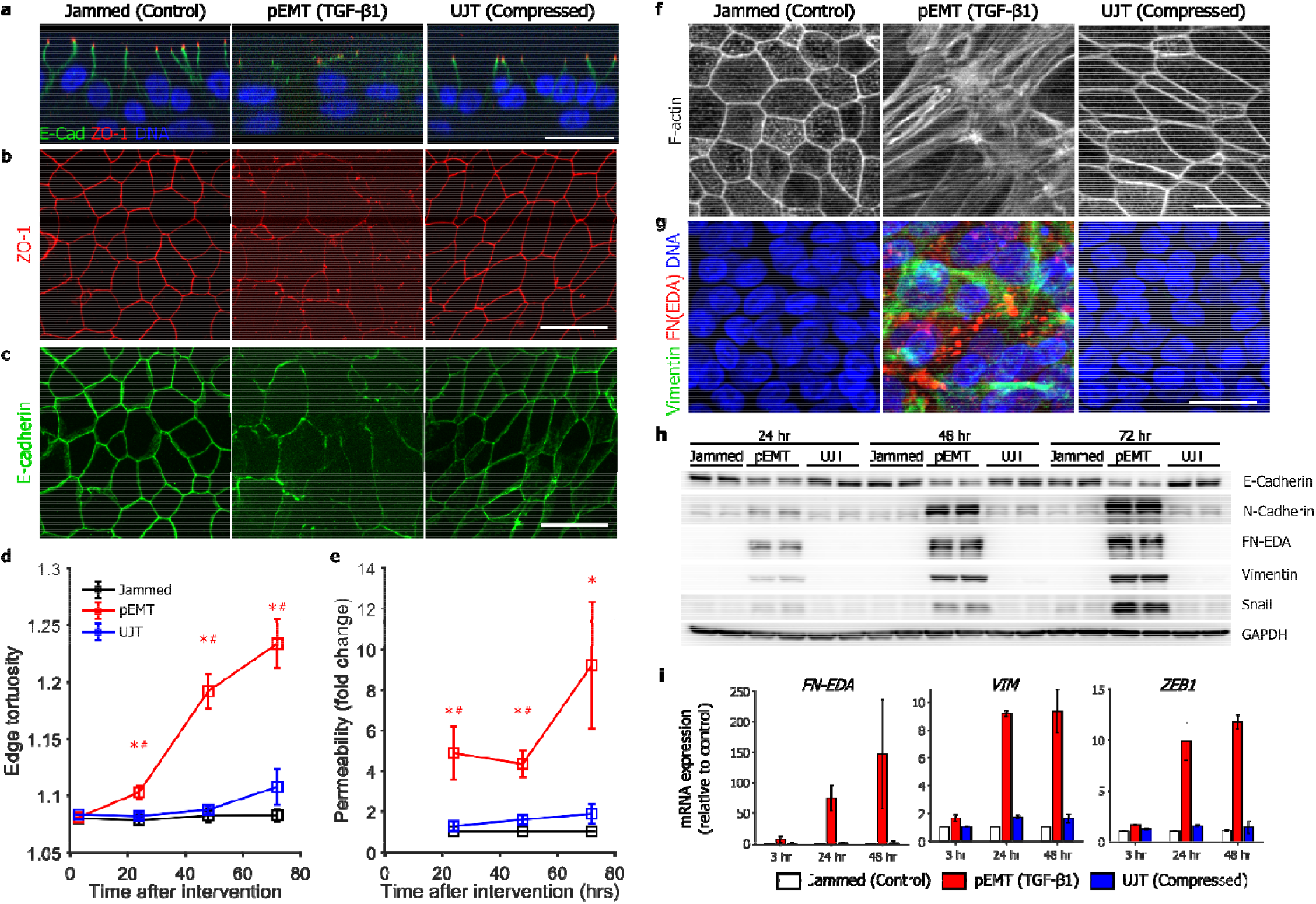
UJT occurs without evidence of pEMT. Representative immunofluorescence (IF) images (**a-c**, **f**, **g**) at 48 hrs after stimulus for jammed (control), pEMT (TGF-β1-treated), and UJT (compressed) layers. **a.** In both jammed and unjammed layers, ZO-1 (red) is localized at the apical tight junctions, while E-cadherin (green) is localized at lateral adherens junctions, consistent with the epithelial phenotype; DNA is shown in blue. In pEMT layers, both ZO-1 and E-cadherin are delocalized from apical and lateral junctions, consistent with the mesenchymal phenotype. **b**, **c.** In b th jammed and unjammed layers, apical tight junctions (ZO-1, **b**) and lateral adherens junctions (E-cadherin, **c**) remain intact, while in pEMT layers the cell edges comprised of these junctions become disrupted. **d.** During UJT, cells elongated while maintaining straight cell edges, while during pEMT, cell-cell edges become progressively tortuous. **e.** Permeability of the layer measured by the dextran-FITC (40 KDa) flux assay was significantly increased during pEMT but remained almost unchanged during UJT (n=4 donors from independent experiments). **f.** During pEMT, cortical actin becomes disrupted while apical stress fibers emerge, indicating loss of epithelial character and gain of mesenchymal character. During UJT, cells maintain intact cortical F-actin; aside from elongated cell shape, cortical actin in jammed versus UJT was indistinguishable. **g.** IF images stained for mesenchymal marker proteins: cellular fibronectin (the Extra Domain A splice variant, denoted FN-EDA, red) and vimentin (green). FN-EDA and vimentin are expressed during pEMT but not during UJT. Vimentin appears as basally located fibers, while FN-EDA appears as cytoplasmic globules. Staining in jammed and unjammed layers were indistinguishable from the isotype control (not shown). **h.** During pEMT, E-cadherin protein levels progressively decreased while mesenchymal markers, N-cadherin, FN-EDA, and vimentin, and the EMT-inducing transcription factor (TF) Snail, progressively increased. During UJT, these protein levels remained unchanged. Western blot quantifications from n=3 donors from independent experiments are shown in Extended data Figs. 1 and 2. **i**. Uring pEMT, mRNA expressions of mesenchymal markers, *VIM* and *FN-EDA* and the EMT-inducing TF *ZEB1*, are significantly elevated, but during UJT remain unchanged. Side view images in **a** were reconstructed from a z-stack, while top down images were maximum intensity projections generated either from the apical-most (**b**, **c**, **f**) or basal-most (**g**) ~10 μm of the z-stack. Scale bars in all images are 20 μm. See also Extended Data Figs 1 and 2. *p<0.05, vs. control; ^#^p<0.05, pEMT vs. UJT, color coded according to which sample is referenced.

### Structure

We segmented cells from images of cell boundaries labeled for E-cadherin or ZO-1 (Extended Data Fig. 1) and quantified cell shapes using the cellular aspect ratio (AR), calculated as the ratio of the major axis to the minor axis of the cellular moment of inertia^11^. Control HBE cells exhibited a cobblestone, rounded, and relatively uniform appearance with 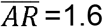 (Fig. 1d, e). Following exposure to compression, however, these cells became more elongated and variable, with progressive growth of 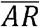 to 2.3 at 72 hrs (Fig. 1d, e, Extended Data Table 1). Following exposure to TGF-β1, cells elongated to 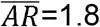 at 24 hrs but plateaued thereafter (Fig. 1d, e, Extended Data Table 1). As discussed below, the boundaries of cells treated with TGF-β1 also exhibited increased edge tortuosity (Fig. 2d).

After compression, both cell migration and elongation grew over time (Fig. 1). In agreement with previously published work, these data indicate that the control unperturbed cells exhibited dynamic and structural signatures of a jammed epithelium, while the compressed cells exhibited dynamics and structural signatures of an unjammed epithelium^9–11^. After TGF-β1 treatment, cell migration and elongation initially increased but migration thereafter tapered off and cell shapes remained unchanged. By these dynamic and structural metrics, UJT and pEMT were indistinguishable at 24hrs but subsequently diverged. Overall, UJT and pEMT showed distinct profiles of cell migration and shape.

## After compression-induced migration, epithelial character persists

We next investigated the extent to which molecular signatures of pEMT and UJT were distinct or overlapping. Control cells exhibited prominent tight junctions marked by apical ZO-1 and adherens junctions marked by lateral E-cadherin (Fig. 2a-c, Extended Data Fig. 1a-c). ZO-1 and E-cadherin appeared as ring-like structures, demarcating continuous cell boundaries and forming cell-cell junctions (Fig. 2b, c). Cell boundaries were relatively straight (Fig. 2d), suggesting that cell-cell junctions were under the influence of mechanical line tension^63, 64^. Further, cells exhibited a cortical F-actin ring, which is a hallmark feature of mature epithelial cells (Fig. 2f, Extended Data Fig. 2a), and they exhibited only undetectable or very low expression of mesenchymal markers (Fig. 2g-I, Extended Data Fig. 2b, c). In well-differentiated, mature pseudostratified HBE cells which are jammed, these data serve as a positive control for fully epithelial character.

Exposure to TGF-β1 (10 ng/ml) disrupted epithelial architecture and led to acquisition of mesenchymal character (Fig. 2, Extended Data Figs. 1 and 2). Importantly, the transition through pEMT to full EMT strongly varies depending on both the degree and the duration of the EMT-initiating signal. As shown in Extended Data Fig. 3, full EMT of well-differentiated HBE cells required extended exposure to TGF-β1; here we focus on pEMT. As expected, in response to TGF-β1, both apico-basal polarity and tight and adherens junctions, as marked by ZO-1 and E-cadherin, became progressively disrupted (Fig. 2a-c, Extended Data Fig. 1a-c), and the level of E-cadherin protein decreased (Fig. 2h, Extended Data Fig. 1d). Remaining cell-cell junctions stained for E-cadherin showed increased tortuosity (i.e., the ratio of edge contour length to edge end-to-end distance) suggestive of a reduction in line tension (Fig. 2d)^63, 64^. Furthermore, to confirm disruption of barrier function, we measured barrier permeability using dextran-FITC (40 kDa)^53^ and observed a substantial increase (Fig. 2e). Cells progressively lost their cortical actin ring while acquiring abundant apical and medial F-actin fibers, a phenotypical feature of mesenchymal cells^65^ (Fig. 2f, Extended Data Fig. 2a). Cells also acquired increased expression of EMT-inducing transcription factors including Zeb1 and Snail1, and mesenchymal markers including N-cadherin, vimentin and fibronectin (EDA isoform) (Fig. 2g-i, Extended Data Fig 2b, c). Increased expression of these mesenchymal markers occurred simultaneously with disruption of epithelial junctions, thus indicating a clear manifestation of a hybrid E/M phenotype and pEMT. In HBE cells undergoing pEMT, these data serve as a positive control for loss of epithelial character and gain of mesenchymal character.

Exposure to compression (30 cm H_2_O) impacted neither apico-basal polarity nor junctional integrity, as indicated by the apical localization of ZO-1 and lateral localization of E-cadherin (Fig. 2a, Extended Data Fig. 2a). These junctions were continuous (Fig. 2b, c, Extended Data Fig. 1b, c) and nearly straight, thus indicating that during UJT the junctional tension was largely maintained (Fig. 2d). Unlike during pEMT, during UJT the overall level of E-cadherin protein remained unaffected (Fig. 2h, Extended Data Fig. 1d). During UJT cells maintained an apical cortical F-actin ring (Fig. 2f, Extended Data Fig. 2a). While barrier function was compromised during pEMT, it remained intact during UJT (Fig. 2e). By contrast to cells during pEMT, cells during UJT did not gain a detectable mesenchymal molecular signature (Fig. 2g-i, Extended Data Fig. 2). These data show that epithelial cells undergoing UJT, in contrast to pEMT, maintained fully epithelial character and did not gain mesenchymal character. Thus, UJT is therefore distinct from EMT.

## During UJT, cellular cooperativity emerges

To further discriminate between pEMT and UJT, we next focused on cooperativity of cell shape orientations and migratory dynamics. Because of immediate cell-cell contact in a confluent collective, changes of shape or position of one cell necessarily impacts shapes and positions of neighboring cells; cooperativity amongst neighboring cells is therefore a hallmark of jamming^39, 40, 46, 66, 67^. We measured cooperativity in two ways. First, we used segmented cell images to measure cell shapes and shape cooperativity that defined structural packs (Fig. 3a). We identified those cells in the collective that shared similar shape-orientation and then used a community-finding algorithm to identify contiguous orientation clusters (methods, Fig. 3a). In both jammed and pEMT layers, cellular collectives formed orientational packs that contained on the order of 5-10 cells and remained constant over time (Fig. 3c). After UJT, by contrast, collectives formed orientational packs that contained 45 ± 22 cells at 24 hrs and grew to 237 ± 45 cells by 72 hrs (mean ± SEM, Fig. 3c, Extended Data Table 1).

**Figure 3.**
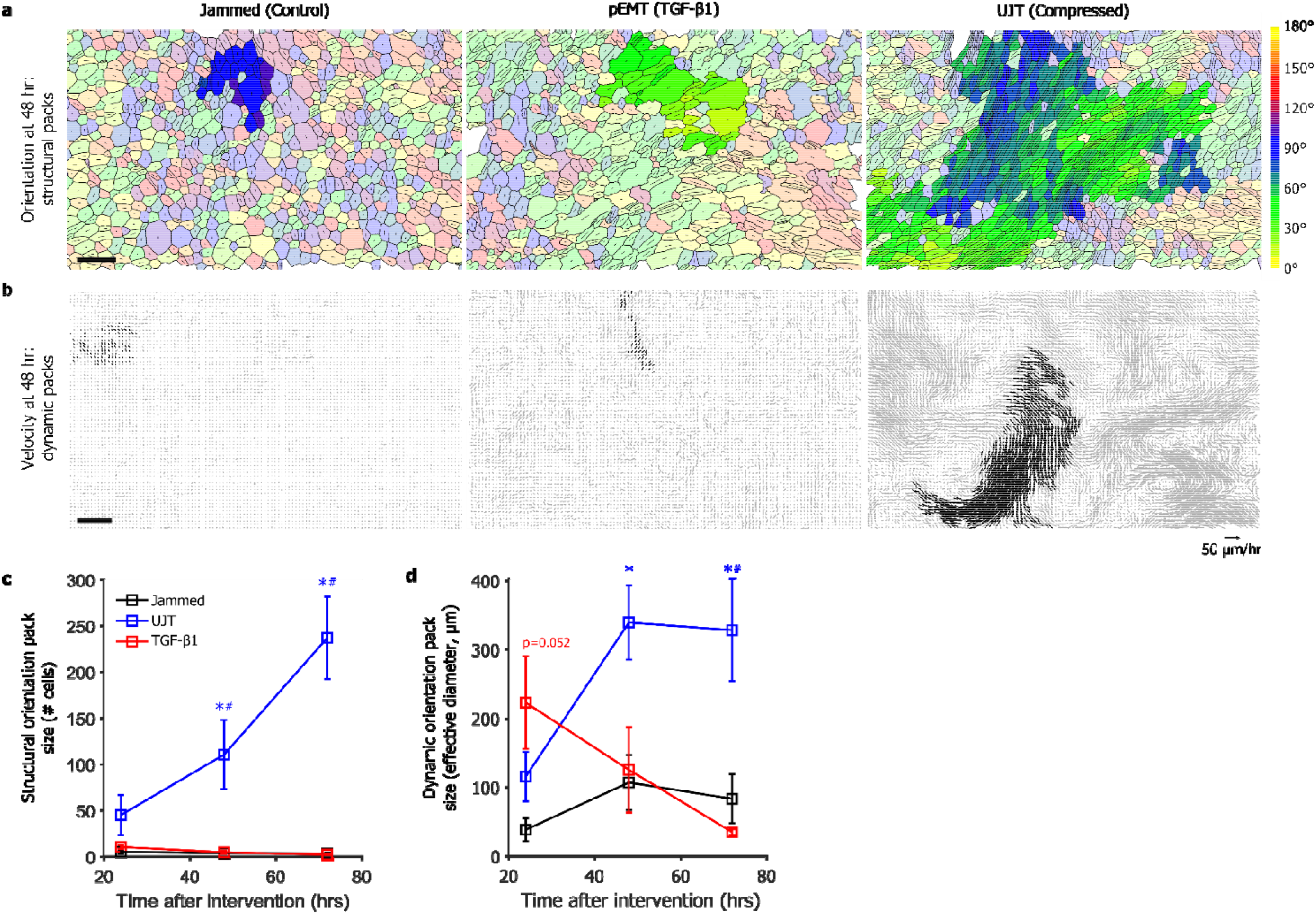
During UJT but not pEMT, cellular alignment and cellular migration organize into cooperative packs. **a.** Using a community-finding algorithm (see methods), cellular orientations were seen to align into orientational packs. Each cell is shown with an orientation director whose length is proportional to AR. Largest orientational packs highlighted in bold colors. Scale bar is 50 μm. **c.** Cells did not exhibit large structural packs when jammed or during pEMT at any time point (n=3 donors from independent experiments). During UJT, by contrast, cells aligned into packs whose median size progressively grew in cell number from 24 to 72 hrs. **b.** Using velocity fields obtained by optical flow and the same community-finding algorithm, cellular motions were seen to organize into oriented migratory packs. Velocity vectors are shown for a 2-hrs of period, with members of the largest dynamic pack highlighted in black. Scale bar is 100 μm. **d**. In jammed layers dynamic packs remained relatively small at all time points (n=4 donors from independent experiments). During pEMT dynamic packs were substantially larger at 24 hrs but returned to baseline by 48 hrs. During UJT, by contrast, dynamic packs grew dramatically in size and remained elevated out to 72 hrs. See also Extended Data Table 1. *p<0.05, vs. control; ^#^p<0.05, pEMT vs. UJT, color coded according to which sample is referenced.

Second, we used cellular trajectories to measure dynamic cooperativity that defined migratory packs (Fig. 3b). Using optical flow over cell-sized neighborhoods^68^, cellular trajectories were constructed by integration. We then used the same community-finding algorithm as above, but here applied to trajectory orientations rather than cell shape orientations (methods, Fig. 3b). As a measure of effective pack diameter we used (4*a*/*π*)^1/2^, where *a* is pack area. In jammed layers, cellular collectives exhibited small dynamic packs spanning 76 ± 31 µm and containing approximately 11 ± 7 cells (methods, Extended Data Table 1). Interestingly, during pEMT, cells initially moved in dynamic packs spanning 223 ± 67 µm containing approximately 71 ± 29 cells at 24 hrs, but these packs disappeared over a time-course matching the disruption of the tight and adherens junctions (Fig. 2b, c, Fig. 3b, d). By contrast, during UJT cellular collectives initially exhibited relatively smaller dynamic packs spanning 115 ± 36 µm containing approximately 19 ± 9 cells at 24 hrs, but grew to packs spanning 328 ± 74 µm containing approximately 139 ± 55 cells at 72 hrs (Fig. 3b, d, Extended Data Table 1).

Cooperativity of cell shape variations and of cell migration were assessed by different methods but nonetheless showed strong mutual concordance. After UJT, but not after pEMT, structure and dynamics became increasingly cooperative. Taken together with Fig. 2, Extended Data Figs. 2 and 3, these observations indicate that coordinated cellular movement during UJT occurred in conjunction with maintenance of epithelial morphology and barrier function (Table 1). These data are consistent with an essential role for intact junctions in cellular cooperation^69–73^, but are the first to show emergence of coordinated cellular migration in a fully confluent epithelium with no evidence of pEMT.

**Table 1.** Across dynamic, structural, and molecular characteristics, pEMT and UJT are distinct. Across all characteristics, pEMT and UJT diverge. Trends are reported for pEMT and UJT, relative to the jammed condition. These findings establish that UJT is sufficient to account for vigorous epithelial layer migration even in the absence of pEMT.

| Evidence | pEMT | UJT |
| --- | --- | --- |
| Motility | ↑ | ↑↑↑ |
| Cell elongation | ↑ | ↑↑↑ |
| Junctional Integrity | ↓ | Intact |
| Layer integrity | Disrupted | Intact |
| Apical/basal polarity | Lost | Intact |
| Epithelial markers | ↓ | Intact |
| Mesenchymal markers | ↑ | Not detected |
| Structural packs | No | Yes; increased over time |
| Cooperative motion | Initial, then lost | Yes; increased over time |

## pEMT versus UJT: discriminating among fluid-like phases

Results above identify two migratory phases, one arising from pEMT and the other from UJT. To better understand these two distinct collective movement phenotypes and to discriminate the mechanical factors that differentiate them, we extend previous theoretical analyses in the class of so-called vertex models^9, 11, 61, 62, 74–77^. In this extended vertex model, referred to here as the Dynamic Vertex Model (DVM), each cell within the confluent epithelial layer adjusts its position and its shape so as to minimize mechanical energy. This energy, in turn, has three main drivers: one that depends on deformability of the cytoplasm and associated changes of cell area; one that depends on extensibility of the apical actin ring and associated changes of its perimeter; and one that depends on homotypic binding of cell-cell adhesion molecules, such as cadherins, together with extensibility of attendant contractile elements and associated changes in cell perimeter^9, 74, 75^. These structures and associated energies, taken together, give rise to a model parameter that is called the preferred cell perimeter, p_0_, and determines the tension borne along the cell-cell junction, here called edge tension^9, 74, 75^. Importantly, contributions of cortical contraction and cell-cell adhesion to system energy are of opposite signs and are therefore seen to be in competition^78^; cortical contraction favors a shorter cell perimeter whereas cell-cell adhesion favors a longer cell perimeter. Equivalently, decreasing cortical contraction causes edge tension to decrease whereas decreasing cell-cell adhesion causes edge tension to increase. As elaborated in Supplement 1, our extended version of the vertex model departs from previous analyses by allowing cell-cell junctions to become curved and tortuous, much as is observed during pEMT. Edge tortuosity can arise in regions where the effects of edge tension becomes small compared with differences in intracellular pressure between adjacent cells.

In the DVM, increasing p_0_ mimics well progressive disruption of the cell-cell-junction and is thus seen to reflect the known physical effects of pEMT (Fig. 4a). For example, when p_0_ is small and propulsive forces are small the cell layer remains jammed (panel i). Cells on average assume disordered but compact polygonal shapes^9, 74^ and cell-cell junctions are straight. As p_0_ is progressively increased cell shapes become progressively more elongated and cell edges become increasingly curvilinear and tortuous, as if slackened (panels ii, iii). Indeed, edge tensions progressively decrease (as depicted by grayscale intensities) with a transition near p_0_ = 4.1, at which point edge tensions approach zero and edge tortuosity begins to rise (Fig. 4b). Loss of edge tension coincides with fluidization of the layer and a small increase in cell speed (inset), at which point the shear modulus^79^ and energy barriers vanish (Extended Data Fig. 4a). Importantly, for p_0_ to increase as cell-cell adhesion diminishes, as necessarily occurs as pEMT progresses, DVM suggests that cortical contraction must diminish even faster. Vanishing edge tension in the fluidized state is consistent with the notion that EMT weakens cell-cell contacts, and junctions therefore become unable to support mechanical forces.

**Figure 4.**
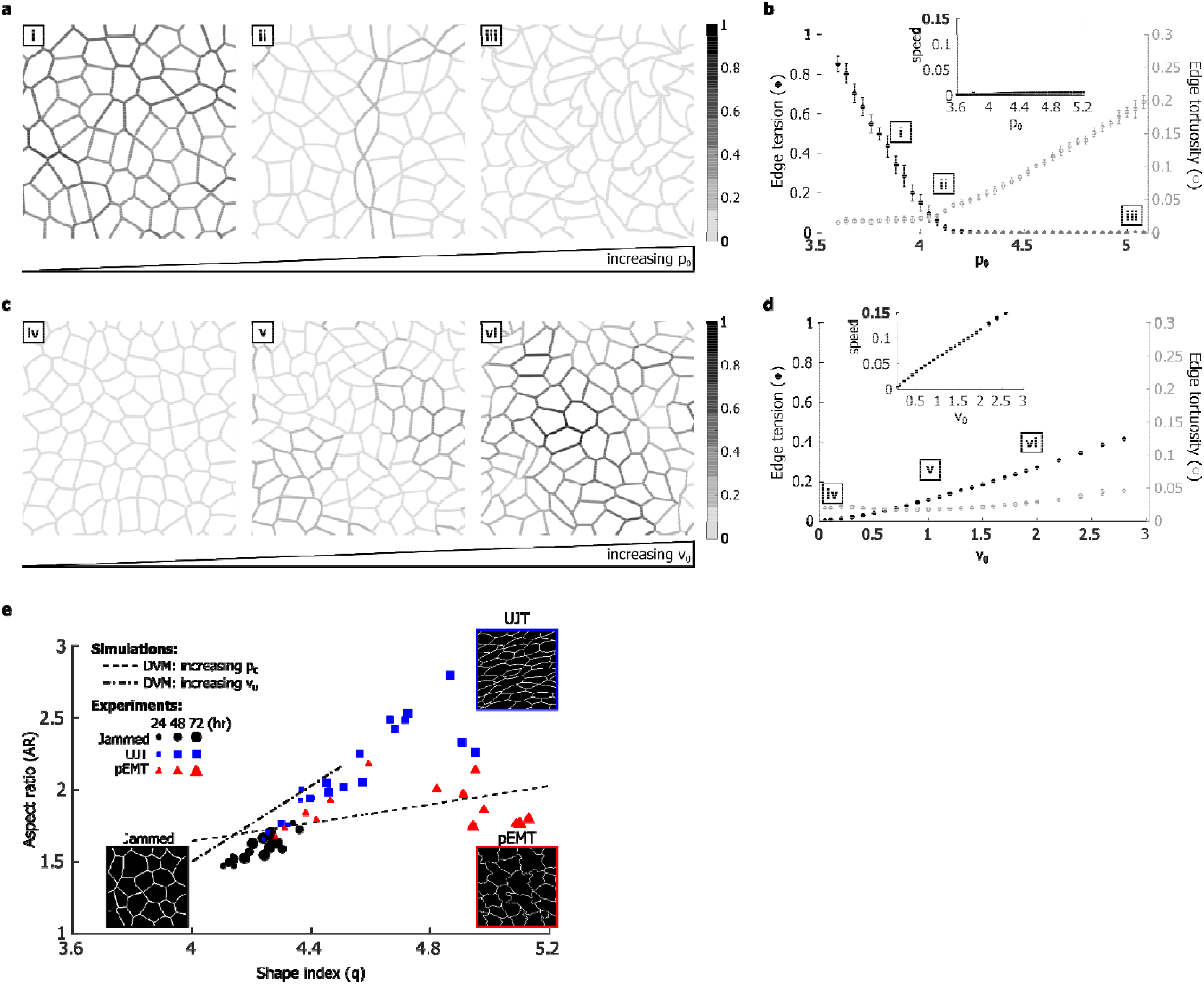
Propulsion, tension, and cell shape discriminate UJT from pEMT. The DVM, an extended vertex model, attributes the effects of pEMT mainly to diminished edge tension but attributes those of UJT mainly to augmented cellular propulsion. **a.** When p_0_ is small and propulsive forces are absent, the cell layer is jammed (i). Cells on average assume compact polygonal shapes^9, 74^, and cell-cell junctions are straight. As p_0_ progressively increases cell shapes become elongated (ii) and cell edges become increasingly curvilinear and tortuous, as if slackened (iii). Further, as p_0_ increases, tension in the cell edges decreases (as depicted by grayscale intensities). **b**. Increasing p_0_ decreased mean edge tension (•), with a transition near p_0_ = 4.1, at which point edge tensions dropped to near zero and edge tortuosity began to rise (○). This loss of edge tension coincides with fluidization of the layer, at which point the shear modulus^79^ and energy barriers vanish (near p_0_ = 4.1; Extended Data Fig. 4a). **c.** If p_0_ is moderate (p_0_=4) and propulsive force v_0_ is small the cell layer is immobile (iv). As propulsive force v_0_ is progressively increased, however, cell shapes become elongated but cell-cell junctions remain quite straight (v, vi). Further, as v_0_ progressively increases, edge tensions increase. **d**. Increasing propulsion v_0_ increases edge tension (•) but without an increase in edge tortuosity (○). Simultaneously, the speed of the cell migration increases (inset). This increase in cell speed coincides with fluidization of the layer: cellular propulsion becomes sufficient to overcome energy barriers that impede cellular rearrangements. **e**. DVM predicts that during UJT versus pEMT two different metrics of cell shape diverge; aspect ratio (AR) emphasizes elongation whereas shape parameter q emphasizes perimeter (q=perimeter/(area^1/2^)). Increasing p_0_ (- - -) moderately increases AR but subs antially increases q, resulting in somewhat elongated cells with tortuous edges. By contrast, increasing 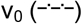 substantially increases AR but minimally increases q, resulting in elongated cells with straight edges. Measurements of AR and q from cells undergoing UJT 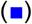 or pEMT 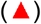 are consistent with those predictions. Theory and observations, taken together, suggest that layer fluidization by means of UJT versus pEMT follow divergent pathways. During pEMT edge tension decreases as junctional adhesion decreases, and as cells elongate q increases more quickly than AR. Cell-cell junctions become increasingly tortuous and slack. During UJT, by contrast, edge tension increases as cellular propulsion v_0_ increases, AR and q increase in tandem, and cells elongate. Cell-cell junctions remain straight and taut.

When propulsive forces, v_0_, are increased while p_0_ is kept fixed, results mimic well the known physical effects of UJT (Fig. 4c). Cell shapes become progressively elongated but cell edges remain straight (panels iv, v, vi). Edge tension increases but without an increase in edge tortuosity (Fig. 4d). Simultaneously, the speed of the cell migration increases appreciably (inset). This increase in cell speed coincides with fluidization of the layer, at which point cellular propulsion has become sufficient to overcome energy barriers that impede cellular rearrangements (Extended Data Fig. 4c).

During UJT versus pEMT, the DVM predicts, further, that two different metrics of cell shape diverge (Fig. 4e). The cellular aspect ratio (AR) emphasizes cellular elongation while deemphasizing tortuosity whereas the shape parameter, q (perimeter/(area^1/2^)) also depends on elongation but emphasizes tortuosity. Measurements of AR versus q from cells undergoing UJT versus pEMT are consistent with those predictions (Fig. 4e). As regards cell shapes and their changes, UJT versus pEMT follow divergent pathways. Together, these results attribute the effects of pEMT mainly to diminished edge tension but attributes those of UJT mainly to augmented cellular propulsion. As such, this new extended vertex model, DVM, provides a physical picture that helps to explain how the manifestations of pEMT and UJT on cell shape and cell migration are distinct.

## Implications

Development, wound repair, and cancer metastasis are fundamental biological processes. In each process cells of epithelial origin are ordinarily sedentary but become highly migratory. To understand the mechanisms by which an epithelial layer can transition from sedentary to migratory behavior, the primary mechanism in many contexts had been thought to require EMT or pEMT^5, 80–83^. During EMT/pEMT cells lose apico-basal polarity and epithelial markers, while they concurrently gain front-to-back polarity and mesenchymal markers. Each cell thereby frees itself from the tethers that bind it to surrounding cells and matrix and assumes a migratory phenotype. In the process, epithelial barrier function becomes compromised. Here by contrast we establish the UJT as a distinct migratory process in which none of these events pertain. Collective epithelial migration can occur through UJT without pEMT or EMT.

EMT/pEMT refers not to a unique biological program but rather to any one of many programs, each with the capacity to confer on epithelial cells an increasingly mesenchymal character^80^. In doing so, EMT/pEMT tends to be a focal event wherein some cue stimulates a single cell ‒or some cell subpopulation– to delaminate from its tissue of origin and thereafter migrate to potentially great distances^25, 84^. As such, EMT likely evolved as a mechanism that allows individual or clusters of epithelial cells to separate from neighbors within the cell layer and thereafter invade and migrate as solitary entities (EMT) or small groups (pEMT) through adjacent tissues^85^. Like EMT/pEMT, the UJT is observed in diverse contexts and may encompass a variety of programs^9–11, 36–44^. But by contrast with EMT/pEMT, UJT comprises an event that is innately collective, wherein some cue stimulates cells constituting an integrated tissue to migrate collectively and cooperatively^86^. Our data suggest that UJT might have evolved as a mechanism that allows epithelial rearrangements, migration, remodeling, plasticity, or development within a tissue under the physiological constraint of preserving tissue continuity, integrity, and barrier function.

We establish here that UJT does not require pEMT, but that finding in turn motivates three new questions. First, UJT has now been observed across diverse biological systems^9–11, 36–44^, but we do not yet know whether UJT is governed across these diverse systems by unifying biological processes or conserved signaling pathways. Second, although we now know that UJT can occur in the absence of pEMT, it remains unclear if pEMT can occur in the absence of UJT. This question is illustrated, for example, by the case of ventral furrow formation during gastrulation in the embryo of *Drosophila melanogaste*r, which requires the actions of EMT transcription factors^87–89^. Prior to full expression of EMT and dissolution of cell-cell junctions in *Drosophila*, embryonic epithelial cells have been shown to unjam; cell shapes elongate and become more variable as cells begin to rearrange and migrate^11^. Supporting that notion, our data in HBE cells point towards a role for UJT in the earliest phase of pEMT; when junctional disruption and expression of EMT transcription factors and mesenchymal markers are apparent but minimal (24 hrs), cells are seen to unjam, elongate, and migrate in large dynamic packs. These observations argue neither for nor against the necessity of EMT for progression of metastatic disease^80, 82, 90, 91^, but do suggest the possibility of an ancillary mechanism.

In many cases the striking distinction between pEMT/EMT versus UJT as observed here is unlikely to be so clear cut. It has been argued, for example, that EMT-induced intermediate cell states are sufficiently rich in their confounding diversity that they cannot be captured along a linear spectrum of phenotypes flanked at its extremes by purely epithelial versus mesenchymal states^5, 16^. In connection with a cellular collective comprising an integrated tissue, observations reported here demonstrate, further, that fluidization and migration of the collective is an even richer process than had been previously appreciated. Mixed epithelial and mesenchymal characteristics, and the interactions between them, are thought to be essential for carcinoma cell invasion and dissemination^16^, but how UJT might fit into this physical picture remains unclear. More broadly, the Human Lung Cell Atlas now points not only to dramatic heterogeneities of airway cells and cell states, but also to strong proximal-to-distal gradients along the airway tree^92^. But we do not yet know how these heterogeneities and their spatial gradients might impact UJT locally, or, conversely, how UJT might impact these gradients. In that light, the third and last question raised by this work is the extent to which pEMT/EMT and UJT might work independently, sequentially, or cooperatively to effect morphogenesis, wound repair, and tissue remodeling, as well as fibrosis, cancer invasion and metastasis^93^.

## Supporting information

Supplement: Model

## Acknowledgments

The authors thank Jeffrey M. Drazen for his critical feedback. The authors acknowledge the support of the Northeastern University Discovery Cluster.

## Author Contributions

J.A.M., M.A.N., J-A.P. designed experiments; J.A.M., M.J.O., I.T.S. performed experiments; A.D. and D.B. performed dynamic vertex model simulations; J.A.M., A.D., M.J.O., S.K., D.B. analyzed data; J.A.M., A.D., S.J.D., J.P.B., J.J.F., M.A.N., D.B., J-A.P. interpreted data; J.A.M., A.D., J.J.F., D.B., J-A.P. wrote the manuscript.

## Methods

### Cell Culture

Primary human bronchial epithelial (HBE) cells at passage 2 were differentiated in air-liquid interface (ALI) as described previously^9, 52, 53, 55, 56^. Briefly, cells were plated onto type I collagen (0.05mg/ml) coated transwell inserts and maintained in a submerged condition for 4-6 days. Once the layer became confluent, media was removed from the apical surface and the ALI condition was initiated. Over 14-17 days in ALI, the cells differentiate and form a pseudostratified epithelium which recapitulates the cellular architecture and constituency of the intact human airway^11, 48, 49, 51, 94^. For the entire culture period, HBE cells were maintained in serum-free media as described in ref ^53^. For the experiment, cells were maintained for 20 hrs with minimal medium depleted of epithelial growth factor, bovine pituitary extract and hydrocortisone. For experiments with time points longer than 24 hrs, cells were fed with fresh minimal media at 48 hrs following the initial media change prior to exposure.

Experiments were repeated with primary cells from at least n=3-4 donors in independent experiments. HBE cells were derived from donors with no history of smoking or respiratory disease, as used in our previous studies^9, 52, 53, 55, 56^. Experimental quantifications are shown across all donors and reported n is number of independent donors used.

To initiate pEMT, cells were treated with recombinant human TGF-β1 (10 ng/ml, Cell Signaling Technology) ^59^. To initiate UJT, cells were exposed to mechanical compression with an apical-to-basal pressure differential of 30 cm H_2_O as described previously^9, 52–56^. Time-matched control cells were set up with vehicle treatment for TGF-β1 and a shame pressure for mechanical compression.

### Protein and mRNA expression analysis

We detected protein levels by western blot analysis as described previously^53^. Cell lysates were collected at 24, 48, and 72 hrs after initial exposure to stimuli (vehicle/sham, TGF-β1 at 10 ng/ml, or compression at 30 cm H_2_O). The following antibodies and dilutions were used: E-cadherin (1:10,000), N-cadherin (1:1000), Snail (1:1000), vimentin (1:1000), GAPDH (1:5000), all from Cell Signaling Technology; EDA-fibronectin (1:1000, Sigma). We report fold-changes of normalized protein levels compared either to vehicle control (for E-cadherin) or to TGF-β1—treated at 72 hrs (for mesenchymal markers) across n=3 donors.

We detected mRNA expression as previously described^56^. Cells were collected from the conditions and donors as described above at 3, 24, and 48 hrs after the initial exposure to stimuli, and RNA was isolated from cell lysates using the RNAeasy Mini Kit (Qiagen) following the manufacturer’s instructions. Real-time RT-PCR was performed using primers listed below, and fold changes were calculated by the comparative ΔΔCt method^95^.

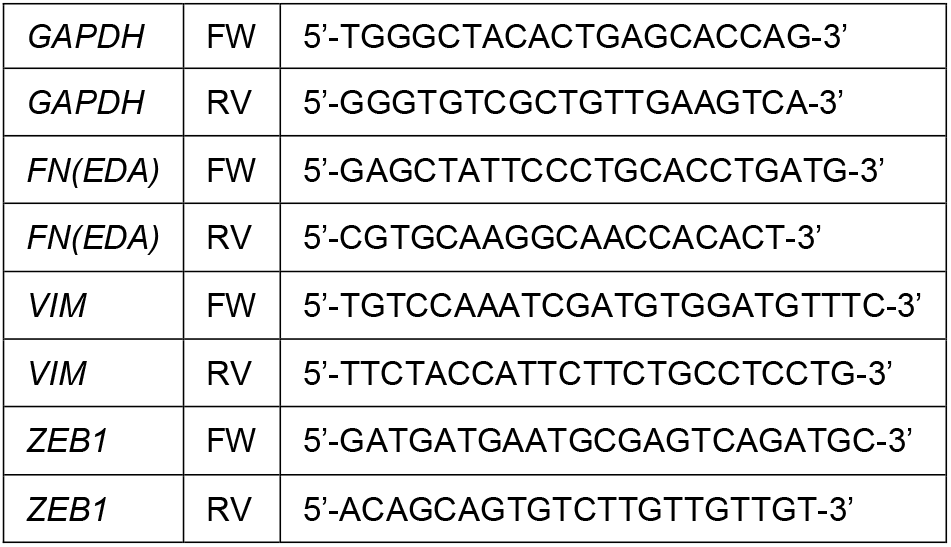
Primers.

### Immunofluorescence

At 24, 48 or 72 hrs after initial exposure to stimuli, cells were fixed with either: 4% paraformaldehyde in PBS with calcium and magnesium for 30 mins at room temperature; or, 100% methanol at −20°C for 20 mins. Cells were permeabilized with 0.2% Triton X-100 for 15 mins and blocked with 1% bovine serum albumin and 10% normal goat serum for 1 hr. Cells were stained for F-actin (Alexa fluor 488-Phalloidin, 1:40, 30 mins) or for proteins of interest, as follows: E-cadherin (1:200, Cell Signaling Technology), ZO-1 (1:100, ThermoFisher), vimentin (1:100, Cell Signaling Technology), cellular fibronectin (1:200, EMD Millipore). Cells were counterstained with Hoechst (1:5000) for nuclei. Following staining, transwell membranes were cut out from the plastic support and mounted on glass slides (Vectashield). Slides were imaged on a Zeiss Axio Observer Z1 using an apotome module. Maximum intensity images were generated in ImageJ.

### Live imaging and dynamic analysis

To determine cellular dynamics, time-lapse movies were acquired and analyzed. Images were taken every 6 min for 6 hrs, ending at 24, 48, or 72 hrs after initial exposure to stimuli. Phase contrast images were acquired on a Zeiss Axio Observer Z1 with stage incubator (37°C, 5% CO_2_). Time-lapse movies were analyzed using custom software written in Matlab. Cellular dynamics were determined using an optical flow algorithm. The movies were registered to sub-pixel resolution using a discrete Fourier transform method^96^. Flow fields were calculated from the registered movies using Matlab’s OpticalFlowFarneback function. Trajectories were seeded from the movie’s first frame using a square grid with spacing comparable to the cell size and obtained from forwards-integration of the flow fields; for our field of view there were about 4000 trajectories. The average speed was calculated from the displacement during a two-hour window, and the effective diffusivity was calculated from the slope of the mean square displacement.

### Permeability

Epithelial barrier function was determined by a dextran-FITC flux assay, as described previously^53^. Directly following time-lapse imaging of HBE cells, 1 mg/ml dextran-FITC (40 kDa; Invitrogen) was added to the apical surface of cells. After 3 hrs, medium was collected from the basal chamber, and was used for measuring fluorescence intensity of FITC. Fluorescence intensity measured in media from stimulated cells is expressed as fold-change relative to that in the media from time-matched control cells.

### Cell shape analysis

To determine cell shape distributions, we marked cellular boundaries and measured shape characteristics as described below. To mark cellular boundaries, we segmented immunofluorescent cell images using SeedWater Segmenter^13^. Images used were maximum intensity projections of ZO-1 and E-cadherin at the apical region of the cell layer. Segmented images were used to determine cell boundaries and extract cell shape information, including apical cell area, perimeter and aspect ratio (AR) from major and minor axes of an equivalent ellipse. This fitted ellipse has equivalent eigenvalues of the second area moment as of the polygon corresponding to the cell boundaries, as published previously^11^. In addition to cell AR, we computed the cell shape index 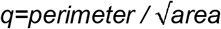. We also extracted individual cell edges and computed the end-to-end distance and the contour distance along the edge to compute the edge tortuosity:

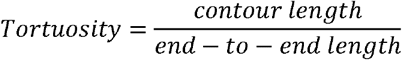

### Structural and dynamic cluster analysis

Orientation clusters, or packs, were determined from both cell shape orientation and from cell trajectory orientation, using a community-finding algorithm as described below. Cell shape orientation was determined from segmented immunofluorescent cell images, while cell trajectory orientation was determined from dynamic flow fields. Each cell or trajectory possessed orientation *θ*_*j*_ with respect to a global axis of reference. The method below was developed for cell shape orientation clusters and was then applied to cell trajectories to determine dynamic orientation clusters. The determination of orientation clusters started by initiating a neighbor-count on each cell in a given image. We detected the number of neighbors *m*_*i*_ of the *i*-th cell possessed similar orientations within a cutoff *δθ* = ±10°. This led to an increase of neighbor-count on each of these neighbor cells of cell *i* by the number *m*_*i*_. We created the set of these neighbor cells for cell *i* and repeated this neighbor-finding for each of the other members in the set except cell *i*. We increased the neighbor-counts on all the members by the newly found number of neighbors and updated the set of connected cells. We continued to look for neighbors for all the new members of the set until we were unable to find a neighbor with similar orientation for any new member. This gave us a cluster of structurally connected cells where each of the cells have at least one neighbor with orientation within *δθ*. We called this an orientation-based cluster or a structural pack. We determined the mean pack-size per cell by counting, for each cell, the number of cells in its pack, and averaging. This can be expressed mathematically as follows: if in the *j*-th structural pack there are *s*_*j*_ number of cells, and there are *N*_*c*_ cells in an image, the mean pack-size per cell would be 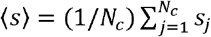. A null test for our algorithm was to set *δθ* = ±90° and find that all cells in an image became part of the same connected cluster giving a mean pack-size equal to the number of cells.

We performed the same pack-size analysis on the cellular trajectories obtained from the velocity field determined using optical flow. We applied a uniform speed threshold equal to the mean speed on each image and then a cutoff on the orientations of velocity vectors given by *δθ* = 10°. The rest of the calculation proceeded as above. Once we obtained the number of velocity vectors in each dynamic pack, we converted this to a two-dimensional area corresponding to the size of the pack. We then expressed an effective pack size according to the (4*a*/*π*)^1/2^, where *a* is pack area. We also converted this areal pack size into an approximate number of cells by using the average cell size determined for control cells for 4 donors, from the shape analysis described above.

### Statistical methods

All of the data was analyzed in Matlab using custom scripts. To determine statistical significance, we ran an ANOVA for each data set, comparing across the multiple donors used. This was followed by a post-hoc analysis using a Bonferroni correction, and p<0.05 was considered significant.

## Data availability

Data that comprise the graphs within this manuscript and other findings of this study are available from the corresponding author upon request.

## EXTENDED DATA

**Extended Data Figure 1:**
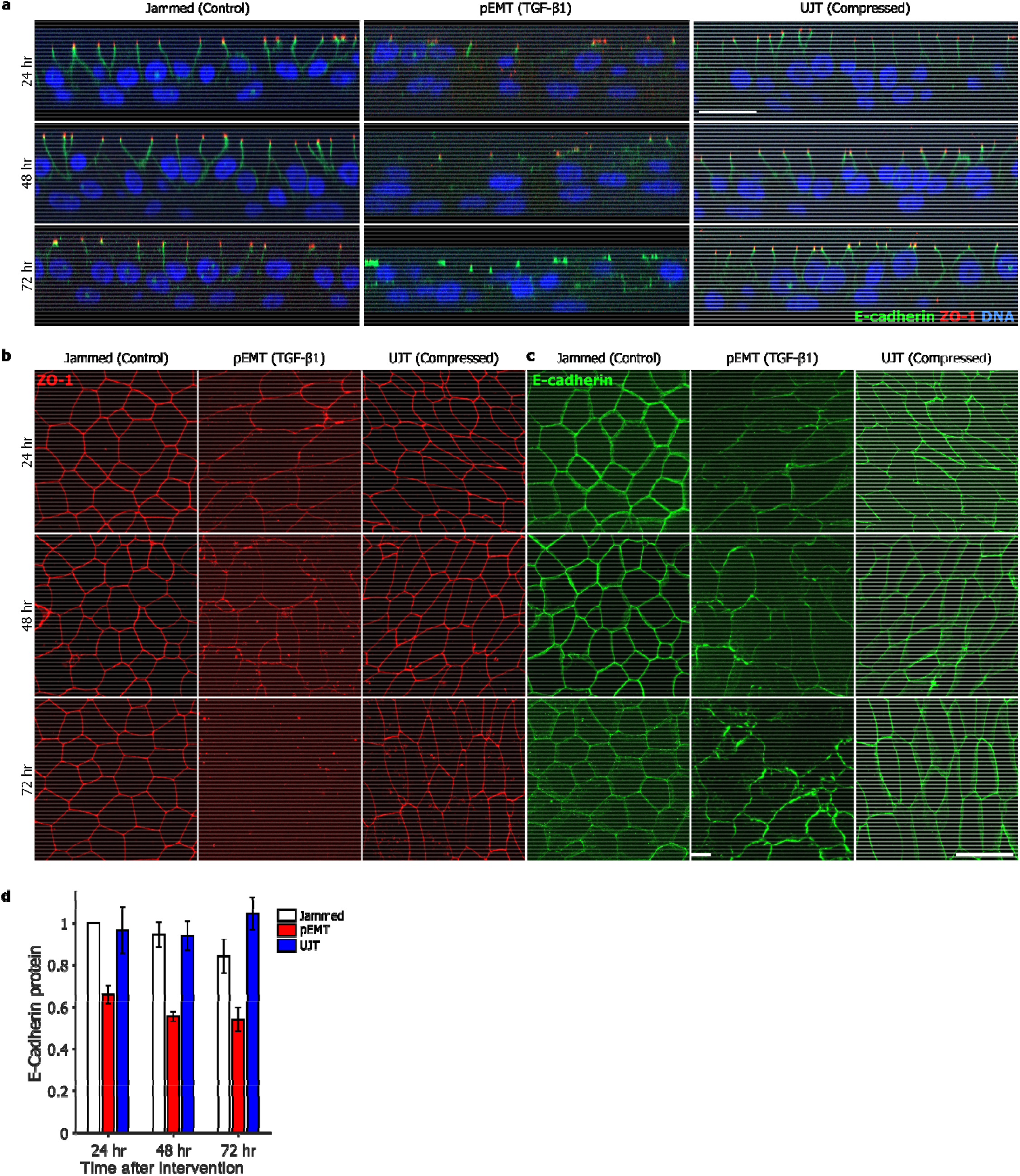
Partial EMT reduces epithelial character, while unjamming maintains epithelial character. Extended data from Figure 2. Representative immunofluorescence (IF) images (**a**-**c**) at 24, 48 or 72hrs after stimulus for jammed (control), pEMT (TGF-β1-treated), and UJT (compressed) layers. **a**, At all time points, in both jammed and unjammed layers, ZO-1 (red) is localized at the apical tight junctions, while E-cadherin (green) is localized at lateral adherens junctions, consistent with the epithelial phenotype; DNA is shown in blue. In pEMT layers, both ZO-1 and E-cadherin are delocalized from apical and lateral junctions, consistent with mesenchymal phenotype. This occurs as early as 24hrs and persists during pEMT. **b**, **c**, At all time points, in both jammed and unjammed layers, apical tight junctions (ZO-1, **b**) and lateral adherens junctions (E-cadherin, **c**) remain intact, while in pEMT layers the cell edges comprised of these junctions become progressively disrupted. **d**, Quantification of protein expression from Fig 2h (n=3 donors). Expression of E-cadherin decreases during pEMT but remains unchanged during UJT.

**Extended Data Figure 2:**
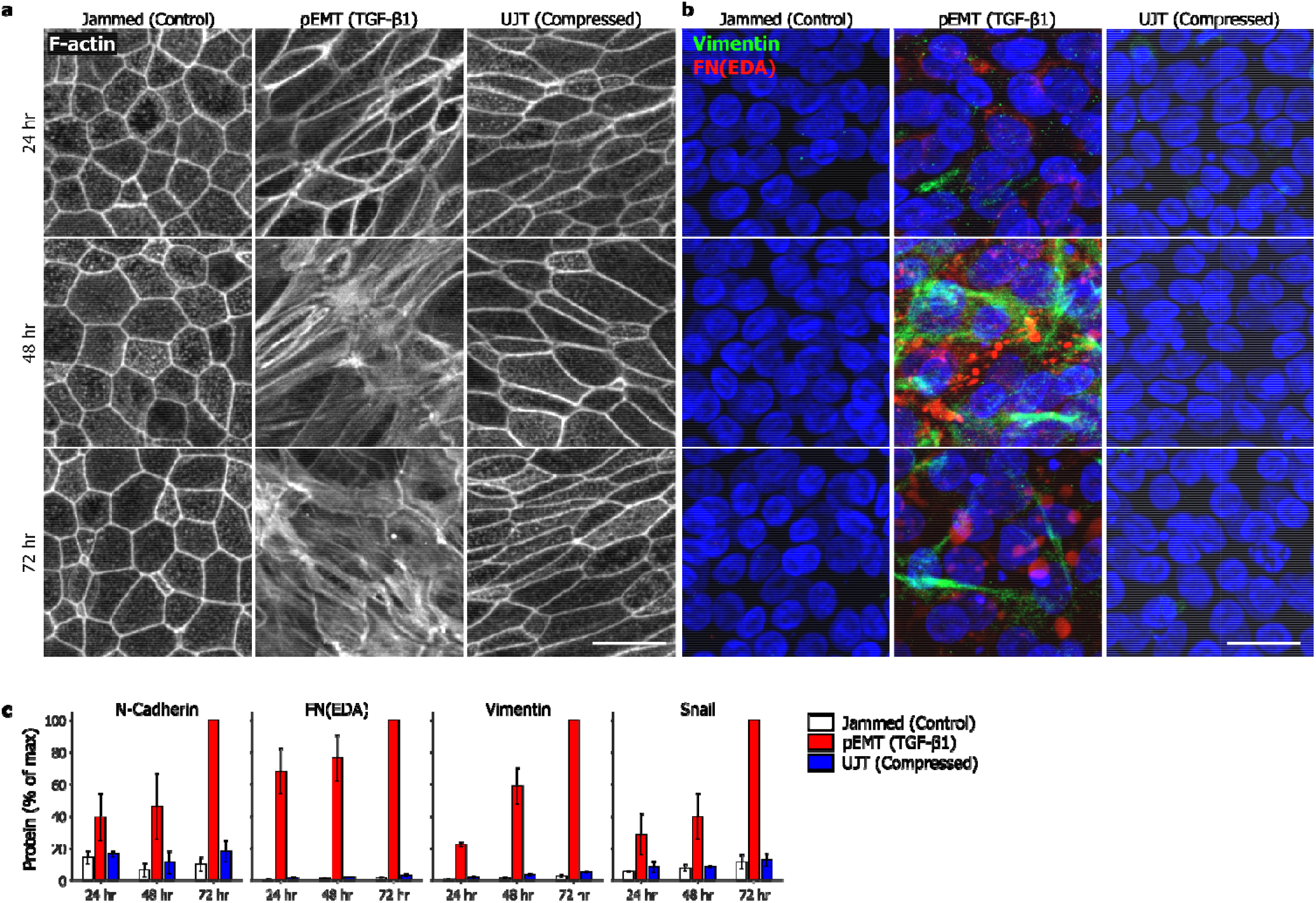
Partial EMT increases mesenchymal character, while unjamming does not. **a**, During pEMT, cortical actin becomes disrupted while apical stress fibers emerge, indicating loss of epithelial character and gain of mesenchymal character (at 48 and 72 hrs). During UJT, cells maintain intact cortical F-actin; aside from elongated cell shape, cortical actin in jammed versus UJT was indistinguishable. **b**, IF images stained for mesenchymal makers: cellular fibronectin (the Extra Domain A splice variant, denoted FN-EDA, red) and vimentin (green). FN-EDA and vimentin are expressed during pEMT but not during UJT. Vimentin appears as basally located fibers, while FN-EDA appears as cytoplasmic globules. Staining in jammed and unjammed layers were indistinguishable from the isotype control (not shown). **c,** Quantification of protein expression from Fig 2h (n=3 donors). During pEMT, mesenchymal markers, N-cadherin, FN-EDA, and vimentin, and the EMT-inducing transcription factor (TF) Snail, progressively increased. During UJT, these protein levels remained unchanged.

**Extended Data Figure 3:**
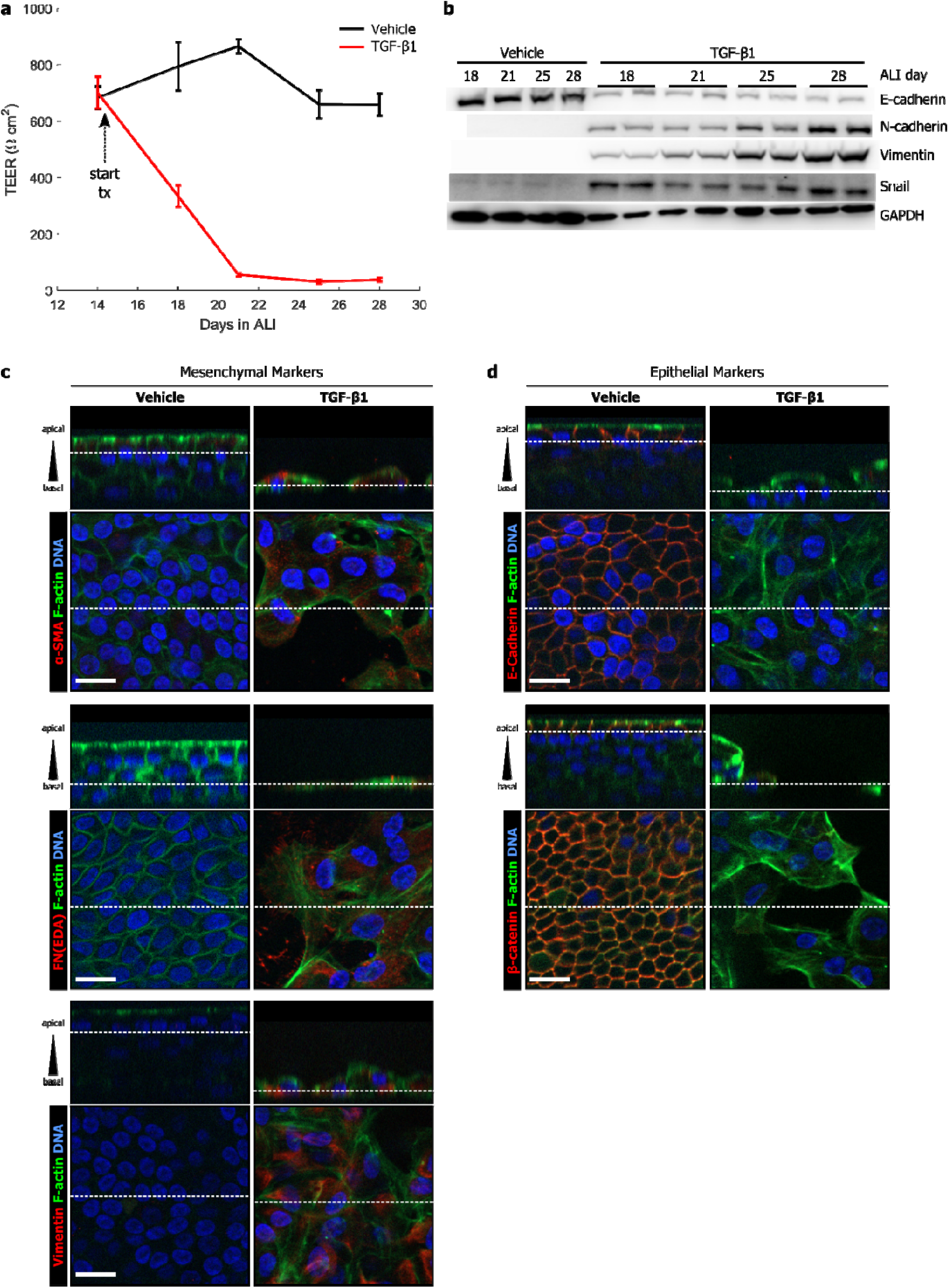
Long exposure to TGF-β1 is required for HBE cells to undergo full EMT. The main text focuses on the initial events during early partial EMT. To elicit a more complete EMT, involving a complete loss of epithelial character and a strong gain of mesenchymal character and cell individualization, we treated differentiated HBE cells with TGF-β1 (10 ng/ml) continuously for up to 14 days, starting on ALI day 14. **a**. Epithelial layer integrity was measured by trans-epithelial electrical resistance (TEER) every four days starting just prior to treatment with TGF-β1 or a vehicle control. By 4 days of treatment, the TEER of TGF-β1—treated HBE cells undergoing EMT was substantially lowered compared to the control cells. By 8 days of treatment, the TEER of the treated cells was negligible, due to large areas of denuded cells and significant breakdown to epithelial junctions. **b**. To determine extent of EMT over the 14-day treatment, epithelial and mesenchymal markers were detected by western blot in HBE cells treated with TGF-β1 or vehicle. Vehicle-treated HBE cells retained expression of E-cadherin and did not acquire expression of N-cadherin, vimentin, or snail. By contrast, TGF-β1—treated HBE cells lost expression of E-cadherin and gained expression of N-cadherin, vimentin, and snail. Expression of N-cadherin and vimentin progressively increased over time. **c**. Immunofluorescence staining of mesenchymal markers (α-SMA, FN-EDA, vimentin) shows that expression of these were increased in response to TGF-β1 treatment (14 days), but that they were not expressed in vehicle-treated control HBE cells. **d**. Immunofluorescence staining of epithelial markers (E-cadherin and β-catenin) shows that expression of these were maintained and localized at the cell-cell junctions in vehicle-treated control HBE cells. In contrast, these epithelial markers were undetectable following TGF-β1 treatment (14 days), where cell contacts were completely disrupted.

**Extended Data Figure 4:**
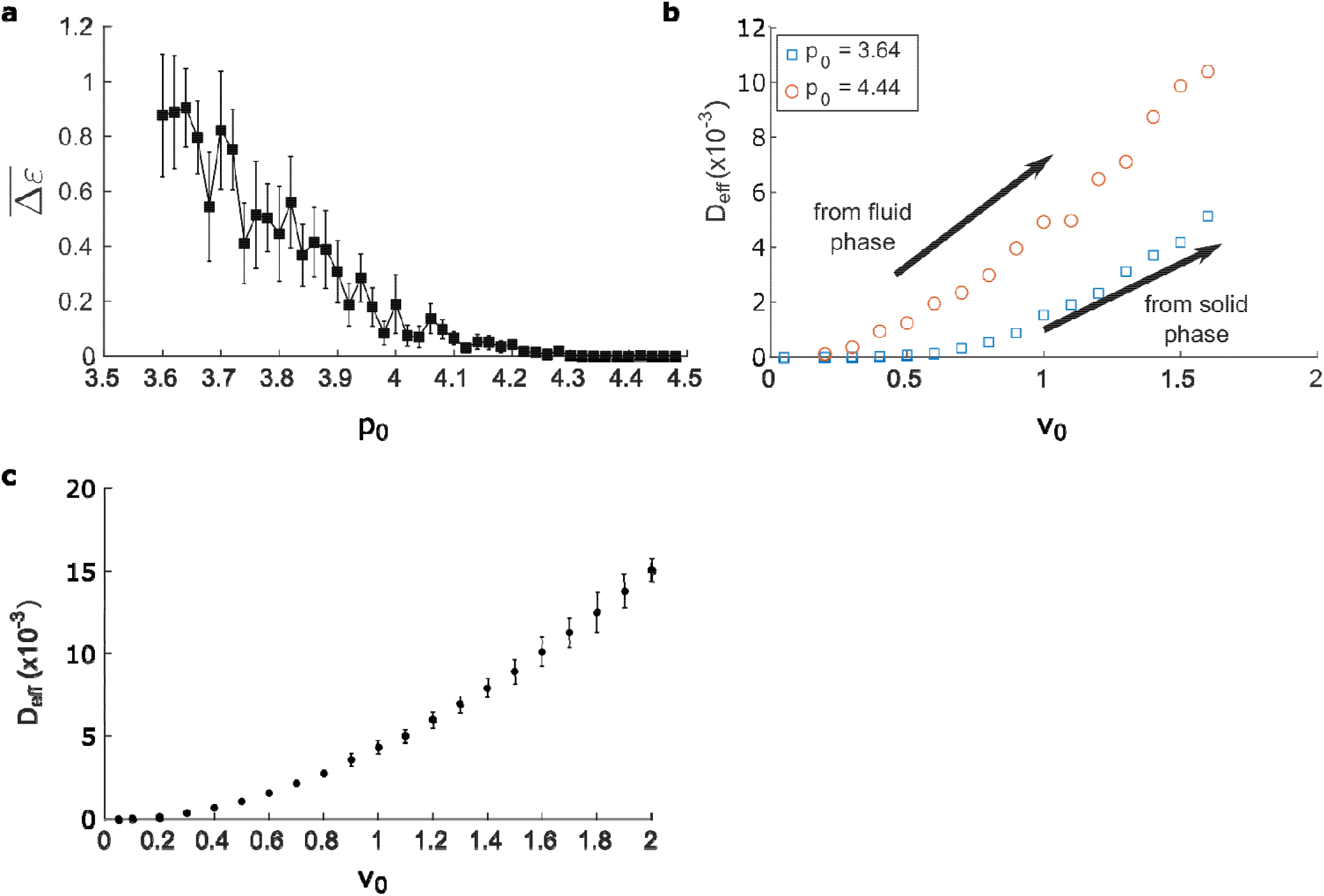
Energy barriers and migration in the dynamic vertex model (DVM). **a,** Average energy barrier to structural rearrangements () for cells in the DVM when p_0_ is increased and v_0_ is small (v_0_=0.05) and cell edges are allowed to curve. Precisely where the edge tensions vanished, near p_0_~4.1, the energy barrier drops to zero. This indicates that cells in this fluid-like phase require only a small increase in v_0_ in order to become motile, as shown in **b**. This mode of migration appears to correspond to pEMT (see Fig 4e). **b,** The effective diffusivity, D_eff_ (defined as) captures the amount of average movement of cells in the DVM and is shown for two representative values of p_0_, as v_0_ is increased. For cells in in the solid-like phase (p_0_=3.64), cellular movement occurs at a substantially higher value of v_0_, compared to cells in the fluid-like phase (p_0_=4.44), which exhibit vanished tension and vanished energy barriers, as in **a**. **c**, When v_0_ is increased and p_0_ is moderate (p_0_=4.0), cells migrated as shown by increasing D_eff_. In this case, cell edges remained under high levels of tension. Though energy barriers to cellular rearrangement were finite (not shown), cellular migration occurred when cellular propulsion was sufficient to overcome these barriers. This mode of migration appears to correspond to the UJT (see Fig 4e).

**Extended Data Table 1.** Biophysical measurements show how pEMT and UJT diverge. Dynamic and structural metrics for HBE cells undergoing pEMT or UJT reported as mean across donors with standard error in parentheses. For dynamic measurements (speed, D_eff_, dynamic pack size), data was obtained from n=4 donors with 6-12 fields of view per time point and condition captured in independent experiments. For structural measurements (AR, structural pack size), data was obtained from n=3 donors with ≥ 2 fields of view per time point and condition captured in independent experiments. Statistics are shown as follows: *p<0.05 vs. control; ^#^p<0.05 UJT vs. pEMT. Statistical significance was determined by an ANOVA followed by post-hoc multiple comparisons tests with Bonferroni correction.

| Condition | Timepoint<br>(hrs after<br>treatment) | Average speed<br>($\mu\text{m/hr}$) | $D_{\text{eff}}$ ($\mu\text{m}^2/\text{hr}$) | Mean aspect<br>ratio (AR) | Structural pack<br>size (#cells) | Dynamic pack<br>size (effective<br>diameter, $\mu\text{m}$) |
| --- | --- | --- | --- | --- | --- | --- |
| Jammed<br>(Control) | 24 | 1.0 (0.4) | 1.6 (1.3) | 1.58 (0.07) | 5 (2.3) | 38 (17) |
|  | 48 | 2.8 (1.2) | 8.3 (3.5) | 1.59 (0.06) | 4 (0.3) | 107 (40) |
|  | 72 | 2.1 (1.0) | 7.9 (4.1) | 1.60 (0.04) | 3 (0.7) | 83 (36) |
| pEMT<br>(TGF-β1) | 24 | 3.0 (0.8) | 6.3 (2.5) | 1.84 (0.08)* | 10 (2.3) | 223 (67) |
|  | 48 | 2.4 (0.4) | 5.0 (1.3) | 1.92 (0.09)* | 4 (1.8) | 125 (62) |
|  | 72 | 1.9 (0.3) | 3.6 (1.1) | 1.83 (0.06) | 2 (0.5) | 35 (4) |
| UJT<br>(Compressed) | 24 | 3.7 (0.9)* | 9.0 (3.0) | 1.82 (0.08)* | 45 (22) | 115 (36) |
|  | 48 | 9.9 (2.0)*<sup>#</sup> | 64.2 (19.7)*<sup>#</sup> | 2.22 (0.12)* | 110 (38)*<sup>#</sup> | 339 (54)* |
|  | 72 | 9.5 (3.0)*<sup>#</sup> | 97.8 (37.6)<sup>#</sup> | 2.30 (0.09)*<sup>#</sup> | 237 (45) *<sup>#</sup> | 328 (74)*<sup>#</sup> |

