## Supplement: Model for "The unjamming transition is distinct from the epithelial-to-mesenchymal transition"

**Supplement 1:
Simulation of collective cellular behavior using an extended Dynamic Vertex Model**

Vertex-based models^1-3^ have been successful in describing cell shape statistics in developing embryos^4-7^ and glassy and jamming behavior in cultured epithelial sheets^8-11^. These computational models are simple yet contain the salient features that offer useful means to investigate the role of cell-cell interactions and how single cell behavior is connected to emergent collective behavior at the multicellular level.

The framework of this model is based on the observation that the cells comprising epithelial sheet tend to maintain a columnar structure and the 2D cross-section of the epithelial sheet forms a polygonal tiling in the plane^1, 2^. The apical surface of cells is then specified by the location of vertices, as well as the geometry of edges that connect the vertices. In this model, the edges represent the conformation of the cell-cell junctions. Unlike classical implementation of the vertex model, here we allow the edges to be curved as an emergent property (as detailed below). The cells in such a tiling emerge as *curved* polygons of different shapes with well-defined area ($A$) and perimeter ($P$).

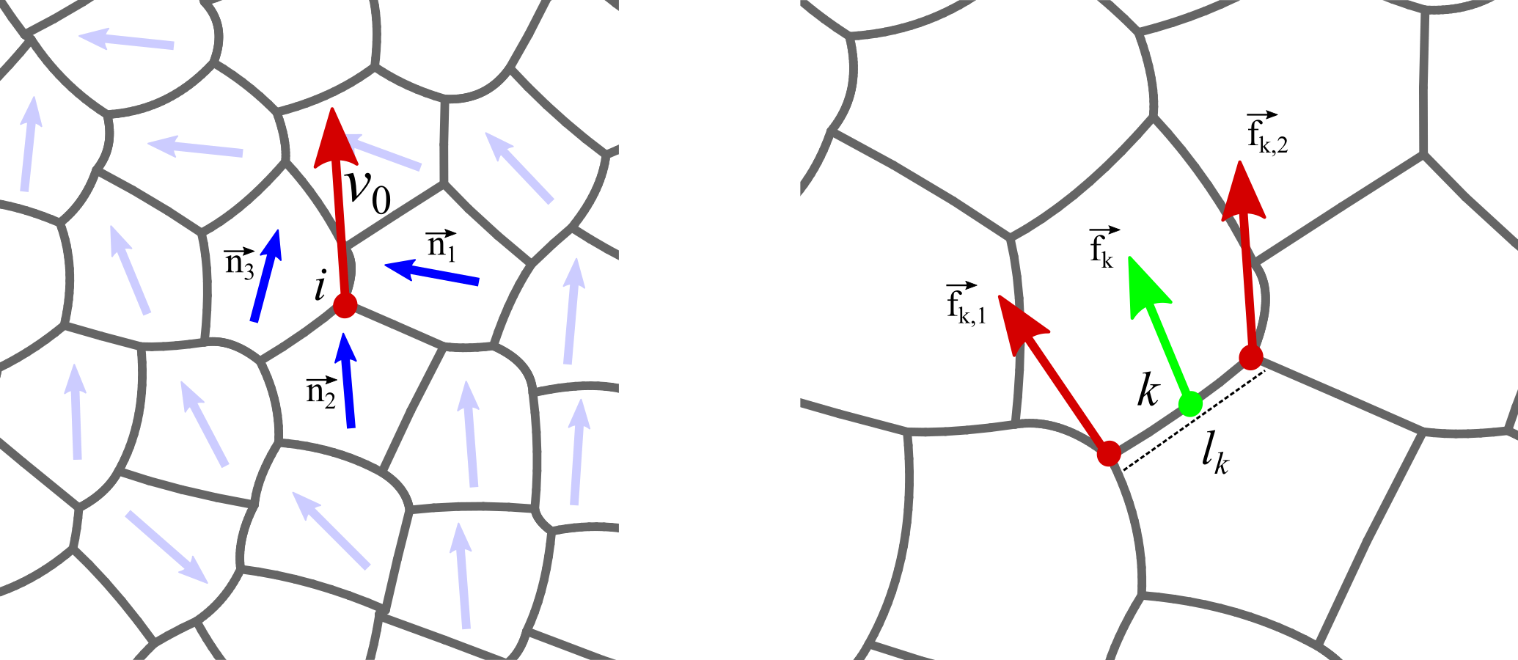

Figure S1. Schematic for the dynamic vertex model which shows a part of the simulated tissue. Left, the net propulsion (solid red arrow, definition Eq. 3) on a given vertex (red dot) has a magnitude $v_{0}$ and direction derived by the polarizations of three cells sharing the vertex, given by $\vec{n_{1}}$, $\vec{n_{2}}$ and $\vec{n_{1}}$ (solid blue arrows). Right, the active force on a mid-edge vertex $k$ given by $\vec{f_{k}}$ (solid green arrow, definition Eq. 5) which depends on the active forces on the head vertex ($\vec{f_{k,1}}$) and tail vertex ($\vec{f_{k,2}}$) (solid red arrows), $l_{k}$ being the end-to-end length of the edge carrying $k$.

The classical vertex model: Beginning with the network of vertices and connecting edges, vertex models then describe the mechanical energy of each cell written as function of cell area (A_i_) and perimeter (P_i_) of the $i$-th cell:

$E_{i}={K_{P}P_{i}}^{2}+\lambda_{i}P_{i}+{K_{A}(A_{i}-A_{0}^{i})}^{2}$	(1)

The first term describes the elastic energy required to deform the cell cortex and the contractile elements embedded within the cell cortex. This term is quadratic in cell-perimeter, $K_{P}{P_{i}}^{2}$ with elastic constant $K_{P}$. The second term describes the energetic effects arising from cell-cell adhesion and cortical contractile tension. This term is linear in cell perimeter, $\lambda_{i}P_{i}$. The coefficient $\lambda_{i}$ can be reduced in two distinct ways ^4, 12^: (1) by increasing the homotypic cadherin bonds on cell-cell junctions or (2) by diminishing cortical contractile tension. Both scenarios would lead to a longer perimeter for the cell^4, 12^.

The first two terms in Eq.1 can be combined into the well-known form^4, 9^: ${K_{P}(P_{i}-P_{0}^{i})}^{2}$ where $P_{0}^{i}=-{\lambda_{i}}/\left( 2K_{P} \right)$, an effective preferred perimeter^13^ of the $i$-th cell. The third term in Eq 1 is associated with the energy cost for changing the cross-sectional area of the$i$-th cell from its rest area $A_{0}^{i}$ with an elastic constant $K_{A}$. This energy arises due to cytoskeletal elasticity and cell deformations that change cell area away from $A_{0}^{i}$ ^5^. Together the perimeter and area energy terms give rise to the vertex model energy function^4, 5, 13^ for $N$ cells:

$E=\sum_{i=1}^{N} \left[ {K_{P}(P_{i}-P_{0})}^{2}+{K_{A}(A_{i}-A_{0})}^{2} \right]$	(1a)

For simplicity, here we treat all the cells as being mechanically identical. We can then rewrite Eq 1a in a dimensionless form as follows,

$\tilde{E}=\sum_{i=1}^{N} \left[ {(p_{i}-p_{0})}^{2}+ {\tilde{K}_{A}(a_{i}-1)}^{2} \right]$	(1b)

wherein $K_{P}A_{0}$ is taken as the natural energy scale and $\sqrt{A_{0}}$ is taken as the natural length scale. The rescaled tissue energy, cell perimeter and cell area now become: $\tilde{E}=E/(K_{P}A_{0})$, $p_{i}={P_{i}}/{\sqrt{A_{0}}}$, $a_{i}={A_{i}}/{A_{0}}$, respectively, and $\tilde{K}_{A}= K_{A}A_{0}/K_{P}$. The rescaled preferred cell perimeter becomes $p_{0}={P_{0}}/{\sqrt{A_{0}}}$ which is also called the target cell shape index^14^. When there is mismatch between the actual length of cell boundary ($p_{i}$) and this preferred value ($p_{0}$), tensions arise on cell edges. If the rescaled perimeters of two cells $i$ and $j$ sharing an edge are $p_{i}$ and $p_{j}$, respectively, then the tension on the shared edge is given by:

$T_{ij}=\left( p_{i}-p_{0} \right)+\left( p_{j}-p_{0} \right)$.	(2)

The Dynamic Vertex Model (DVM): To describe cellular migration, the rate of change of position of the $i$-th vertex, $\vec{r}_{i}$ , at time $t$ is represented by the overdamped equation of motion,

$\Gamma\frac{d\vec{r}_{i}}{dt}=\vec{F}_{i}^{int}+v_{0}\sum_{j\leftrightarrow i} w_{j}\hat{n}_{j}$	(3)

where, $\Gamma$ is describes frictional damping. $\vec{F}_{i}^{\mathrm{int}}$ describes those forces that arise due to cell-cell interactions, and set by a spatial gradient in tissue mechanical energy: $-{\partial E}/{\partial\vec{r}_{i}}$. The last term represents an active motility force $\vec{f_{i}}$ on vertex $i$ which has a magnitude $v_{0}$. Its direction is set by a weighted average of the polarization vectors of cells adjacent to vertex $i$, unlike a flat sum over the associated cell polarizations as described before^8, 15, 16^. The weight factor $w_{j}$for cell *j* (polarization $\hat{n}_{j}$) is given by: $w_{j}={l_{j}}/{{(2Z\Sigma}_{j\leftrightarrow i}}l_{j})$ where $l_{j}$ is the sum of the lengths of the two edges shared by cell $j$ and vertex $i$, $Z$ is the connectivity of vertex $i$ and the division by 2 is done to avoid double counting. This ensures that the active force on any vertex gets the largest contribution from the neighboring cell that contains the longest edges connected to the vertex. We choose the cell polarization vectors in spirit of recent models of self-propelled particles^17-22^: $\hat{n}_{j}=\left( \cos\alpha_{j}, \sin\alpha_{j} \right)$ where the angle of polarization $\alpha_{j}$ follows over-damped dynamics according to:

$\frac{d\alpha_{j}}{dt}=\eta_{j}, \left\langle\eta_{j}\left( t \right)\eta_{k}\left( t' \right) \right\rangle= 2D_{R}\delta_{jk}\delta\left( t-t' \right).$	(4)

This description models the front-back polarity that drives motility in migrating cells^14, 23, 24^. Each polarization vector is subjected to a Gaussian random noise $\eta_{j}$ with zero mean and variance $2D_{R}$. This noise introduces a persistence to the cell-polarization with a period $1/D_{R}$ which we keep unchanged in all our simulations. The force generation on any vertex is shown schematically in Fig. S1.

Curving cell-cell boundaries: In this expanded version of the DVM, we now allow each edge to curve, as, as occurs during pEMT, by introducing a mid-edge vertex which provides an additional degree of freedom on each edge. Like the other vertices, each mid-edge vertex is subjected to active forces derived from the cell motility forces on the neighboring vertices. For the mid-edge vertex $k$ on a given edge, the active force $\vec{f}_{k}$ is calculated using the active forces on the head and tail vertices of the edge using the following:

$\vec{f}_{k}=\frac{1}{2}\left[ \left( 1-\frac{l_{1}}{l_{k}} \right)\vec{f}_{k,1}+\left( 1-\frac{l_{2}}{l_{k}} \right)\vec{f}_{k,2} \right]$	(5)

where $\vec{r}_{k}$ is the position vector of the mid-edge vertex, $\vec{f}_{k,1}$, $\vec{r}_{k,1}$ and $\vec{f}_{k,2}$, $\vec{r}_{k,2}$ are the active forces and position vectors of the head and tail vertices, respectively (Fig. S1). Also, $l_{1}=\left| \vec{r}_{k}-\vec{r}_{k,1} \right|$, $l_{2}=\left| \vec{r}_{k}-\vec{r}_{k,2} \right|$ represent the Euclidian distances of the head and tail vertices from the mid-edge vertex respectively, and $l_{k}=l_{1}+l_{2}$, essentially the end-to-end length of the edge.

We estimate the contour length of an edge from an arc that represents the edge. The parametric equation for any point on the arc representing an edge is given by:

$\vec{R}\left( s \right)= \vec{r}_{k,1}s{(s-1)}/2+ \vec{r}_{k}\left( 1-s \right)\left( 1+s \right)+\vec{r}_{k,2}s{(s+1)}/2$	(6)

where $-1\leq s\leq1$. This is implemented in the surface evolver program^25^. Because we keep the mean cell area fixed and cell-to-cell variations of area are small, pressure differences across cell edges are negligible. Therefore, when $p_{0}$is small, the tissue is deep in the solid state, and the edges are under high tension and hence straight. However, for large $p_{0}$ the edge tension diminishes, and edges curve to accommodate the large perimeter.

Simulation details: We initialized each simulation using independent Voronoi tessellation of randomly placed points posing as cell centers. This gives rise to a confluent polygonal tiling that is random. Then we minimized tissue energy (Eq 1b) to find a state with energy close to the ground state energy at zero motility ($v_{0}=0$) using the conjugate gradient protocol with respect to the vertex locations. Next, we allow finite motility i.e. $v_{0}>0$ and performed the dynamical simulations. The vertex locations are updated by the Euler method using Eq. 3 with a timestep of ${\Delta t= 10}^{-2}\tau$ where $\tau=\Gamma/{K_{P}}$, the unit of time in the DVM. All lengths in DVM was measured in unit of $\sqrt{A_{0}}$ where we use $A_{0}=\bar{A}$, the mean area of the cells which is maintained at unity throughout any simulation. The rotational noise on the direction of cell polarizations is given by Eq. 4 where $D_{R}=1/\tau$ , kept fixed throughout the study.

We determined the effects of independently varying the magnitude of cellular motility force, $v_{0}$ and the preferred cell perimeter, $p_{0}$. When keeping all other parameters fixed, we found that increasing only $p_{0}$ recapitulated observations made during pEMT, while increasing only $v_{0}$ recapitulated observations made during UJT. Note that each of our simulations depends on a single set of $v_{0}$ and $p_{0}$ values which do not change during the simulation. Moreover, for each set of $v_{0}$ and $p_{0}$ values we ran 20 different simulations from independent initial configurations. Thus, all the error bars associated with data from the simulations (Fig. 4 and Fig. S4) are SOM, generated from these 20 different independent trajectories.

From the 20 simulations corresponding to each set of $v_{0}$ and $p_{0}$ values, we calculated the following average quantities across the cells in the simulated tissue: edge tension (Eq. 2), edge tortuosity (Eq. 5), aspect ratio, cell shape index^8, 14^ (defined for cell $i$ as $q_{i}={P_{i}}/{\sqrt{A_{i}}}= {p_{i}}/{\sqrt{a_{i}}}$), velocities and effective diffusivities. The aspect ratio of any cell in the tissue was calculated from the eigenvalues of the shape tensor^10^, generated using the positions of the vertices of the cell.

In a confluent epithelial sheet the basic mode of migration is T1 transitions^26^ in which cells swap positions with their local neigbors^27, 28^. In the DVM we allow T1 rearrangements by implementing an embargo timer on each cell given by parameter $\tau_{T1}$. This sets a lower limit on the time between successive T1s involving a cell. We chose appropriate values for $\tau_{T1}$ in different simulation scenarios. For simulations with $v_{0}>0.05$ (associated with UJT) we use $\tau_{T1}=\tau$, while we use a much larger value $\tau_{T1}={10}^{3}\tau$ for the simulations at $v_{0}=0.05$ (associated with pEMT where T1 processes were rarely observed in the experiments). The threshold length for T1 edge swap was fixed at $l_{c}=0.1$ in simulation length unit. We used periodic boundary conditions in both x and y directions on our simulation box containing $N=400$ cells. All our simulations were implemented using the Surface-Evolver program^25^.
